# A streamlined nanopore-compatible 5PSeq protocol for rapid phenotypic antimicrobial sensitivity testing

**DOI:** 10.1101/2025.11.12.687992

**Authors:** Honglian Liu, Susanne Huch, Ryan Hull, Fabricio Romero Garcia, Lilit Nersisyan, Xiushan Yin, Wei-Hua Chen, Juan Du, Vicent Pelechano

## Abstract

Antimicrobial resistance (AMR) poses a significant threat to public health. Rapid and accurate antimicrobial sensitivity testing is essential to guide effective treatment. Here, we present “simplified 5PSeq” (s5PSeq), a streamlined protocol for profiling 5’ monophosphorylated (5’P) mRNA degradation intermediates that reflect ribosome dynamics *in vivo*. By capturing antibiotic-induced, context-specific ribosome stalling events, s5PSeq provides a molecular proxy for bacterial growth inhibition—offering a molecular phenotypic readout without the need for culturing. s5PSeq reduces library preparation time to under four hours and incorporates a novel rRNA blocking strategy. We demonstrated its clinical utility by identifying erythromycin-resistant and sensitive *Clostridioides difficile* clinical isolates. Combining s5PSeq with real-time nanopore sequencing enables fast AMR diagnosis with as few as 3000 reads. In addition to simplifying the study of 5’P co-translational mRNA decay, our work suggests that utilizing information-rich phenotypic molecular readouts can significantly improve AMR diagnostics.

**Highlights:**

- s5PSeq is a streamlined protocol for profiling 5’P mRNA degradation intermediates.
- Context-specific ribosome stalls can be used to assess phenotypic antimicrobial sensitivity at the molecular level.
- Blocking rRNA sequencing at the ligation step streamlines library preparation and lowers costs and hands-on time.
- Integration with nanopore sequencing allows same-day antimicrobial sensitivity testing in species with 5’-3’ exonuclease.

## Introduction

Antimicrobial resistance (AMR) is a major threat to global health, with estimates suggesting it could cause up to 40 million deaths by 2050 if untreated^1^. Rapid and accurate diagnostics are essential to guide effective treatment and limit the spread of resistant pathogens. However, current antimicrobial sensitivity testing methods rely primarily on culturing-based techniques to assess bacterial phenotypic susceptibility to antibiotics^2^. Culturing approaches remain the gold standard as they directly measure the impact of the antibiotic treatment on growth. However, these methods are time-consuming, and labour-intensive, and limited by the specific growth requirements of different bacteria. For example, AMR detection in *Clostridioides difficile*, a pathogen causing serious diarrhoea, presents significant challenges due to its anaerobic growth conditions and slow proliferation (doubling time 40-70min)^3^. Molecular methods such as PCR allow for the direct detection of AMR-related genes^4^. While in principle faster, molecular approaches do not measure the impact of the antibiotic treatment on cell growth, as the mere presence of resistance genes does not always correlate with phenotypic resistance^5^.

To bridge this gap, we developed a sequencing-based method that provides a molecular phenotypic readout of AMR by capturing the immediate translational response of bacteria to antibiotic exposure without the need for cell division. Specifically, we leverage the presence of 5’ monophosphorylated (5’P) mRNA degradation intermediates, which reflect ribosome dynamics in vivo^6–9^. We have previously demonstrated that mRNA degradation signatures can provide information about ribosome dynamics and co-translational mRNA decay in eukaryotes^6,7^ and bacteria^9,10^. In bacterial species that possess the 5’-3’ RNA exonuclease RNase J, this exonuclease tracks the last translating ribosome, trimming the accessible mRNA in such a manner that the position of the exposed 5’P mRNA indicates the ribosome’s position^9^. Our research has demonstrated that analyzing these mRNA degradation fragments can uncover translational changes in response to environmental stresses and provide insights into ribosome stalls associated with amino acid limitation or context-specific antibiotic stalls^9,10^. Based on that work, we hypothesize that quantifying context-specific ribosome stalls after antibiotic treatment could be utilized to assess phenotypic antimicrobial sensitivity at the molecular level.

Here, we present simplified 5PSeq (s5PSeq), a streamlined protocol optimized for clinical implementation. s5PSeq reduces hands-on time, cost, and complexity while maintaining our ability to quantify high-quality 5’P and ribosome stalling profiles. First, we confirmed that short-term (10 minutes) context-specific ribosome stall is correlated with long-term growth inhibition after antibiotic treatment. We optimised s5PSeq for multiple bacterial species and benchmarked against our previously developed HT-5PSeq^11^. To demonstrate the clinical potential of s5PSeq, we evaluated our ability to identify erythromycin-resistant and sensitive *C. difficile* clinical isolates. We further demonstrate s5PSeq integration with nanopore sequencing, showing that it delivers phenotypic susceptibility profiles within 6-10 hours following RNA extraction from as few as 3,000 reads. This integration enables a rapid and cost-effective assessment of phenotypic antimicrobial resistance within a working shift, with potential for broad application across diverse pathogens and clinical settings.

### mRNA degradation signatures provide insights into bacterial growth following antibiotic treatment

Bacterial cells adjust to environmental changes by modifying their gene expression and altering their cell growth. This response is evident during antibiotic treatment, where variations in cell growth are typically used as indicators of antibiotic effectiveness^12,13^. Building on our previous work showing that ribosomes often stall in a drug- and sequence-specific manner^9,10^, we hypothesized that measuring 5’P mRNA degradation signatures could predict bacterial growth after antibiotic treatment. This approach may be particularly relevant for ribosome-targeting antibiotics in bacteria with RNase J, where ribosome stalls could be more easily connected to the drug’s mechanism of action. To test this hypothesis, we monitored the growth of *Bacillus subtilis* and *Lactiplantibacillus plantarum* under varying concentrations of chloramphenicol and erythromycin, using antibiotic concentrations commonly used for phenotypic antimicrobial susceptibility tests (Figure 1)^14–16^. In parallel, we extracted RNA 10 minutes post-antibiotic exposure and analysed the 5’P mRNA degradation signatures using HT-5PSeq^11^. We observed a clear dose-dependent signature where the extent of antibiotic context-specific ribosome stall^17,18^ was associated with differences in long term growth (Figure 1). In response to chloramphenicol, we observed a clear ribosome stall when alanine is positioned in the E site. This context-specific ribosome stall protects mRNA from co-translational decay by RNase J, resulting in the accumulation of 5’P mRNA degradation intermediates 8 nucleotides upstream of the alanine codon^9^ (Figures 1A and 1C). On the other hand, erythromycin binds to the 50S subunit of bacterial ribosomes, blocking the exit tunnel and preventing peptide chain elongation^19^. It also causes context-specific ribosome stalls at proline-rich sequences and the R/K-x-R/K motif^9,20,21^, that can also be observed by 5PSeq (Figures 1B and 1D). The correlation between the differential degree of ribosome stalling and growth suggests that monitoring mRNA degradation patterns could serve as an effective method for predicting long-term phenotypic responses. However, our results also showed that our latest 5PSeq implementation (HT-5PSeq)^11^, which requires around 9 hours for library preparation, remains too time-consuming for its streamlined use in clinical microbiology labs where results would be required within a working shift.

**Figure 1.**
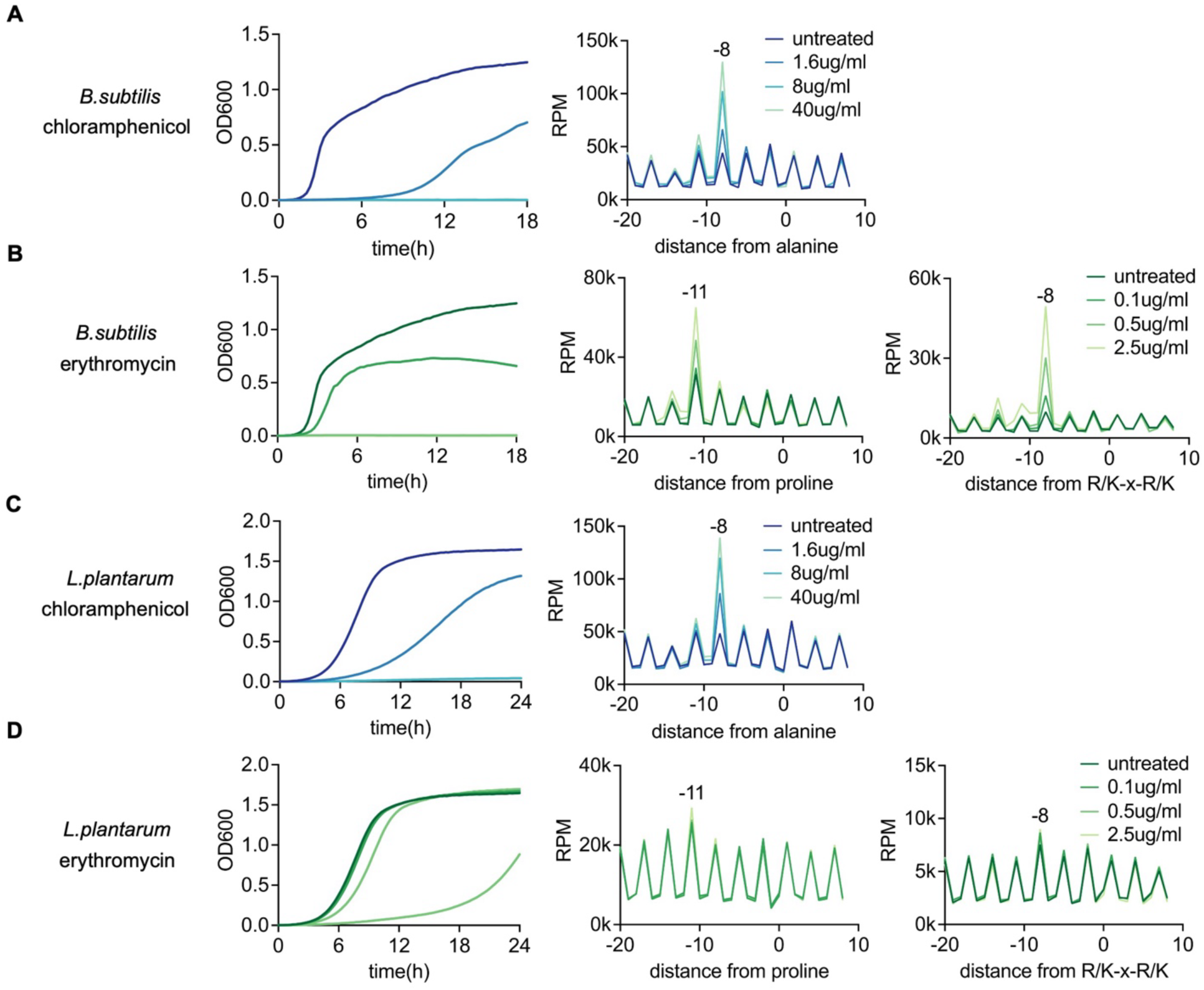
Correlation between long-term growth response and context-specific ribosome stalling after drug treatment. (A) Growth curve (left) and metagene-level context-specific ribosome stalling measured by HT-5PSeq (right) of *B. subtilis* after 10 min of chloramphenicol treatment. The degree of antibiotic context-specific ribosomes stall at position 8nt upstream from alanine is associated with differences in long-term growth. (B) Growth curve (left) and metagene-level context-specific ribosome stalling measured by HT-5PSeq (middle and right) for *B. subtilis* after 10 min of erythromycin treatment. The degree of antibiotic context-specific ribosomes stall measured by 5PSeq at positions 11nt upstream from proline and 8nt upstream from R/K-x-R/K motif (the first base of the last codon of the motif is positioned at 0) is associated with differences in long-term growth. (C) Growth curve and metagene-level context-specific ribosome stalling for *L.plantarum* after 10 min of chloramphenicol treatment, as in (A). (D) Growth curve and metagene-level context-specific ribosome stalling for *L.plantarum* after 10 min of erythromycin treatment, as in (B).

### s5PSeq is a streamlined protocol for profiling bacterial mRNA degradome

To streamline 5’P mRNA degradome sequencing, we have developed a simplified 5PSeq method (s5PSeq), which reduces the library preparation time, pipetting steps, and costs (from 20 USD per HT-5PSeq^11^ library to 10 USD per s5PSeq library using commercial reagents, or 2 USD if using custom-made reagents^22^) while maintaining the quality of the produced sequencing libraries (Figure 2A, STAR Methods and Supplementary Protocol). s5PSeq can be done in under 4 hours and involves rRNA blocking, single-stranded RNA ligation, reverse transcription, and PCR amplification (See Figure S1 for detailed comparison with HT-5PSeq). A key aspect of s5PSeq is a novel approach to prevent abundant 5’P rRNA from dominating the sequencing libraries. Depleting rRNA is crucial when targeting 5’P mRNA, as mature rRNAs dominate the 5’P RNA content in cells. Reducing 5’P rRNA reads allows focusing on mRNA degradation, lowering sequencing costs and enhancing 5PSeq sensitivity. Unlike previous rRNA depletion strategies using biotinylated probes^23^ or targeted degradation of rRNA-containing cDNA molecules^11^, s5PSeq prevents the capture of the 5’P ends of rRNA molecules during the ligation step. Specifically, we utilize a set of short unmodified DNA oligonucleotides (35–40nt) that are reverse complementary to rRNA sequences (Table S1). These oligonucleotides hybridize with rRNAs and effectively block the single-stranded RNA ligation used to identify the 5’P mRNA molecules present in the sample. To ensure that this initial set of rRNA blocking oligos was compatible with a wide range of species, we designed them to block the ligation of rRNA from 10 bacterial species, including model organisms (*Bacillus subtilis*), human pathogens (*Clostridioides difficile, Enterococcus faecalis, Staphylococcus aureus, Streptococcus agalactiae*), and other species commonly found in the human gut, including members of different taxonomic classes such as *Lactiplantibacillus plantarum* (Bacilli), *Roseburia intestinalis*, *Ruminococcus gnavus*, *Dialister invisus*, and *Faecalibacterium prausnitzii* (various classes within Bacillota and other phyla). To enhance the range of rRNAs targeted by the designed probes, in addition to selecting relatively diverse bacteria for our initial set, we incorporated conserved sequences for universally conserved regions, degenerate bases for consensus sequences complementary to less conserved regions, and species-specific sequences for variable regions (Figure 2B).

**Figure 2.**
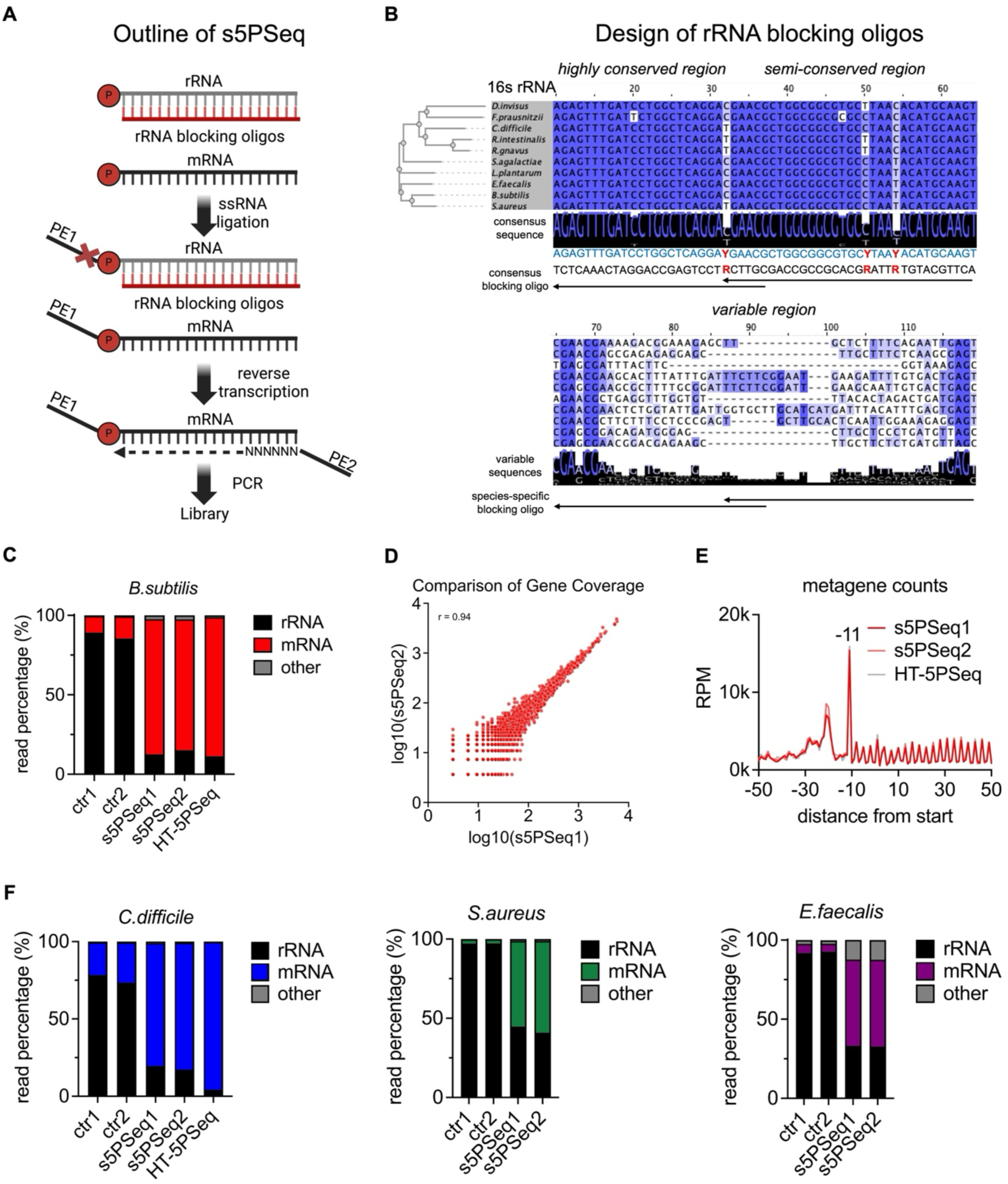
Development of the s5Pseq method. (A) Outline of the s5Pseq protocol. Single stranded DNA oligos are hybridized to total ribosomal RNA to inhibit single-stranded ligation. Following ligation, samples undergo reverse transcription and PCR amplification to produce sequencing-compatible libraries. (B) Design of rRNA blocking oligos based on a multiple sequence alignment of 16S rRNA from 10 bacterial species. Blocking oligos are designed as reverse complements of highly conserved regions, incorporate degenerate bases (e.g., Y for C or T) in semi-conserved regions, or species-specific for variable regions. (C) Relative abundance of reads mapping to rRNA, mRNA and others (tRNAs, ncRNA, scRNA…) obtained using s5Pseq (2 replicates shown) and HT-5Pseq in *B. subtilis.* Control samples (ctr1 and ctr2) omit the rRNA blocking step in s5PSeq. (D) Gene coverage comparison between replicates s5PSeq1 and s5PSeq2 in *B.subtilis*, showing a strong linear correlation (Pearson’s *r* = 0.94). Gene counts were normalized using median ratio normalization, followed by log₁₀ transformation for visualization. (E) Metagene analysis of 5’P mRNA read coverage relative to the start codon in *B. subtilis*. (F) Relative abundance of reads mapped to rRNA, mRNA and others for *C.difficile*, *S.aureus* and *E.faecalis,* as in (C).

We initially evaluated s5PSeq in *B. subtilis* and confirmed that this novel strategy for blocking rRNA ligation achieved comparable rRNA depletion to that provided by HT-5PSeq (88% of mRNA derived reads in HT-5PSeq vs 84% in s5PSeq) (Figure 2C). s5PSeq is highly reproducible when measuring 5’P mRNA degradation abundance per gene (Pearson’s *r* 0.94, Figure 2D) and comparable with HT-5PSeq (Figure S2A). Notably, s5PSeq effectively revealed the anticipated ribosome-associated 5’P mRNA degradation profiles at the translation start and stop, closely resembling those of HT-5PSeq (Figures 2E, S2B and S2C). Finally, s5PSeq was also able to identify expected amino acid specific ribosome stalls (Figures S2D and S2E and Supplementary file for interactive FivePSeq reports^8^) We subsequently tested s5PSeq in clinically relevant bacterial species including *C. difficile*, *S. aureus* and *E. faecalis*. In all cases, s5PSeq provided sufficient coverage of the 5’P mRNA degradome to analyse co-translational mRNA decay (81%, 56%, and 55% mRNA reads respectively, Figure 2F), showed good reproducibility (Figures S2F and S2G) and ribosome-associated 5’P mRNA degradation profiles compared to HT-5PSeq (Figure S2H and S2I). This confirms that s5PSeq can provide high-quality 5’P mRNA degradome information with less hands-on time and cost than previous methods.

### s5PSeq enables detection of phenotypic AMR in clinical *Clostridioides difficile* isolates

Next, we investigated whether s5PSeq could be used to accelerate antimicrobial sensitivity testing (AST) and differentiate between sensitive and resistant bacterial strains. We focused on *C.difficile*, a significant clinical concern due to its role in severe gastrointestinal infections and high recurrence rates^24^. Diagnosing antimicrobial resistance is essential for effective treatment of *C. difficile* infections, and clinical laboratories routinely monitor multidrug resistance of clinical isolates, including resistance to erythromycin^25^. However, conventional phenotypic AST remains challenging due to its strict anaerobic growth requirements, slow growth rate, and the absence of standardized, rapid testing methods^26,27^. These limitations highlight the urgent need for a rapid AST approach that can reliably assess their phenotypic antimicrobial resistance.

We focused on the identification of differential response to erythromycin treatment that leads to differential context-specific ribosome stalls that can be easily measured by HT-5PSeq (Figures 1B and 1D). We tested two *C.difficile* clinical isolates, one erythromycin-sensitive and one erythromycin-resistant, both characterized for phenotypic antimicrobial resistance and tested for growth with erythromycin in our conditions (Figure S3). To ensure that cells were in a metabolically active state, we exposed exponentially growing cell cultures to a clinical epidemiological cut-off value (ECOFF) of erythromycin (1ug/ml) for 10 minutes before RNA extraction. To assess the reproducibility of our approach, we tested 4 independent cultures per strain and performed 5 technical replicates per sample, generating a total of 40 s5PSeq libraries. In the erythromycin-sensitive strain, we readily identified context-specific ribosome stalls at proline-rich sequence and R/K-x-R/K motifs, whereas these stalling events were absent in the resistant strain (Figure 3A). To simplify data interpretation, we transformed the context-specific ribosome stall profiles into a simple metric allowing us to distinguish sensitive and resistant strains (Figure 3B).

**Figure 3.**
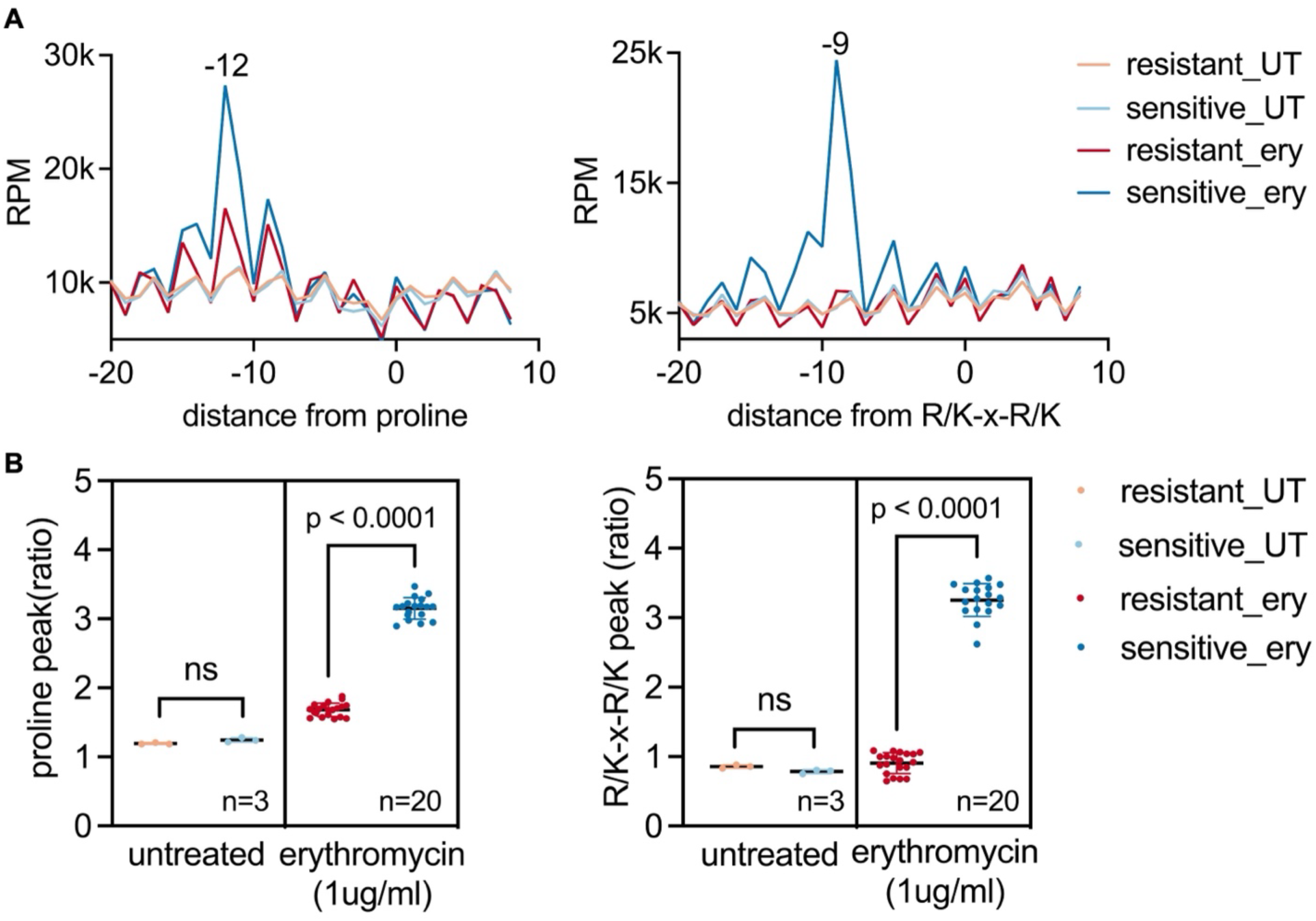
s5PSeq distinguishes erythromycin-resistant and sensitive *C .difficile* clinical isolates. (A) Line plot showing amino acid-specific ribosome stalls measured by 5PSeq at positions 12nt upstream from proline and 9nt upstream from R/K-x-R/K motif (the first base of the last codon of the motif is positioned at 0), sequenced on Illumina NextSeq2000. Context-specific ribosomal pausing is observed exclusively in the sensitive strain under erythromycin treatment ^28^ and not in the untreated samples (UT). (B) Context-specific metric for individual resistant and sensitive samples sequenced from (A). The ratio metric is calculated as the intensity of the 5’P associated to the context-specific ribosomal stall divided by the average intensity of the upstream and downstream 5Pseq coverage. The sensitive strain exhibits a significantly higher peak compared to the resistant strain, as determined by an unpaired t-test (p-val < 0.0001). Only samples with a minimum of 10,000 reads mapped to the coding regions (CDS) were considered.

### Nanopore sequencing coupled with s5PSeq simplifies antimicrobial sensitivity testing

Finally, to further simplify the utilization of s5PSeq as a molecular readout of antimicrobial sensitivity in a clinical context, we examined the potential to integrate it with nanopore-based sequencing approaches. Although short-read technologies (e.g. Illumina) are well-established and offer high-quality reads, their implementation in clinical microbiological settings remains limited due to high costs and their logistical requirements. The high throughput nature of short-read technologies often requires pooling many samples to be cost-effective. Additionally, samples are often transported to an offsite facility, which will delay antimicrobial sensitivity testing and extend the time needed to provide appropriate treatment to patients. Conversely, technologies such as Nanopore sequencing present a more affordable entry cost and, due to their simplified operation requirements, smaller footprint and price, can be easily utilized in clinical microbiology laboratories. Moreover, Nanopore sequencing provides real-time sequencing capabilities and reduces costs related to library preparation and flow cell usage.

We processed the Illumina-compatible s5PSeq PCR amplicons previously prepared for *C. difficile* (Figure 3) by ligating nanopore-specific adapters (SQK-LSK114, 1 hour). We also developed a preprocessing pipeline for nanopore data, including base calling with Guppy/Dorado, demultiplexing, adapter trimming, and read mapping using Minimap2 (see STAR Methods). Due to the relatively higher error rates in nanopore sequencing compared to Illumina, we added a quality control step to filter out low-quality reads by applying read length and quality score thresholds, ensuring only high-confidence reads were used for analysis. First, we sequenced those samples using the PromethION platform known to provide higher quality reads^29^ and throughput. Reassuringly, we were able to clearly identify erythromycin context-specific ribosome stalls (Figure 4A) and distinguish between sensitive and resistant strains (Figure 4B). This was even possible when subsampling reads to 30,000 reads per sample (Figure S4). This suggests that potentially we could multiplex more than 2,000 samples per flow cell in a PromethION run while being able to distinguish sensitive and resistant strains.

**Figure 4.**
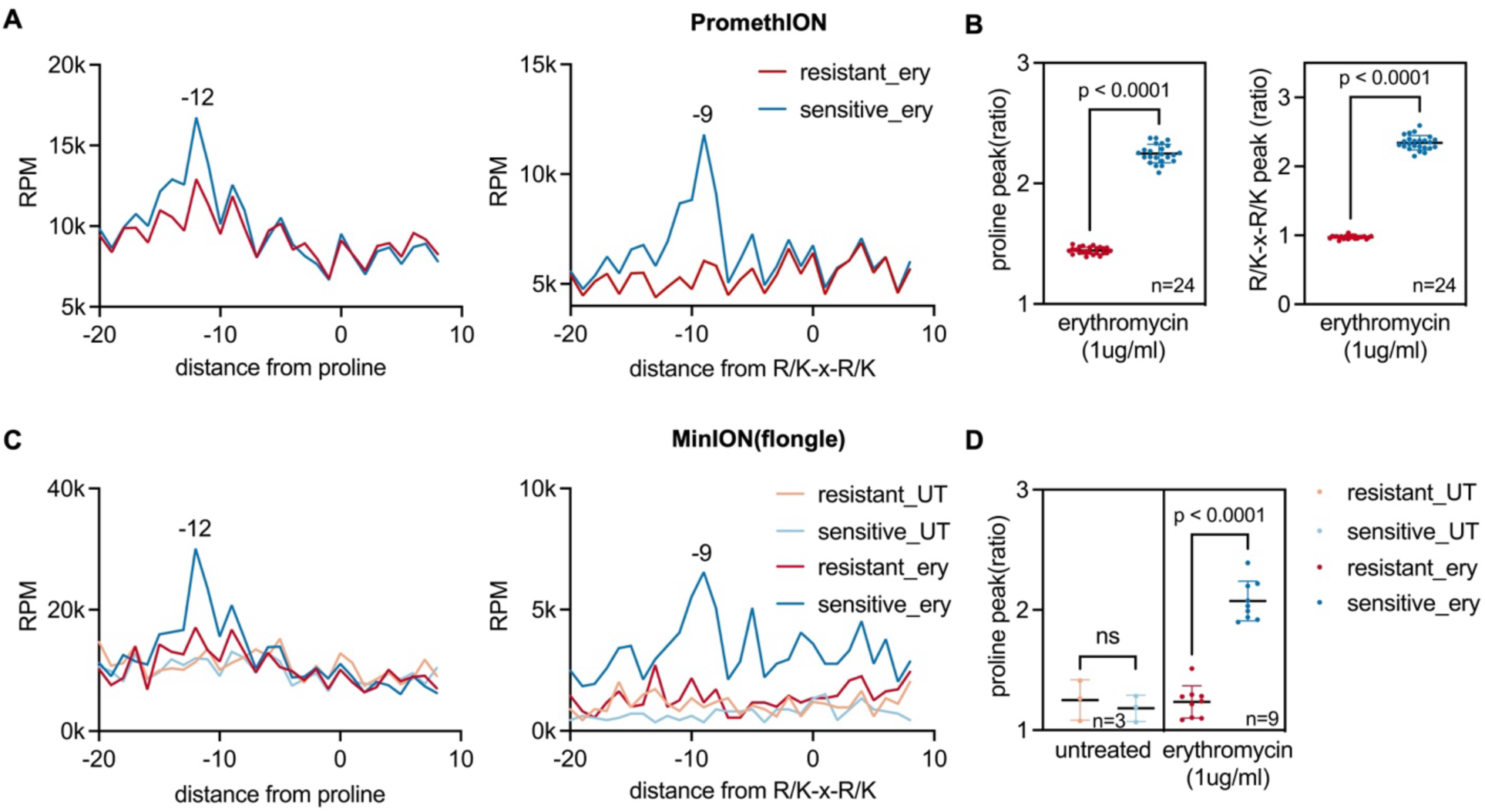
s5PSeq coupled with nanopore sequencing distinguishes erythromycin-resistant and sensitive *C.difficile* clinical isolates. (A) Line plots showing erythromycin context-specific ribosomal stalls at positions 12nt upstream from proline and 9nt upstream from R/K-x-R/K motif (the first base of the last codon of the motif is positioned at 0) measured by s5PSeq and sequenced in a PromethION, as in Figure 3. Replicates are merged yielding 28M (sensitive) and 35M (resistant) reads. (B) Context-specific metric (as in Figure 3B) for individual resistant and sensitive samples sequenced from (A). The sensitive strain exhibits a significantly higher peak compared to the resistant one, as determined by an unpaired t-test (p-val < 0.0001). Only samples with a minimum of 100,000 reads mapped to the coding regions (CDS) were considered. (C) Line plots for s5PSeq libraries sequences in Flongle, as in (A). Replicates are merged and subsampled 30,000 reads. (D) Context-specific metric for individual samples from (C). The sensitive strain exhibits a significantly higher peak compared to the resistant one, as determined by an unpaired t-test (p-val < 0.0001). Only samples with a minimum of 3,000 reads mapped to the coding regions (CDS) were considered.

To further evaluate our ability to use Nanopore sequencing to bring s5PSeq closer to the clinic, we employed the cost-effective ONT Flongle system (priced at 67 USD per flow cell). We sequenced 24 samples per Flongle using the MinION platform. Within five hours, the sequencing generated around 400,000 filtered reads (approximately 10,000 reads per sample after demultiplexing). Despite the relatively low number of reads per sample and the relatively lower sequencing quality associated to the MinION platform when running a Flongle flow cell (average read quality of Phred 8 versus 17 for the same libraries run in a PromethION), the results were sufficient to accurately differentiate between sensitive and resistant strains (Figures 4C and 4D). We could differentiate between strains using as few as 3,000 reads per sample (Figure 4D). Those differences were even more clear when technical replicates where merged and subsampled to 30,000 reads per sample (Figure 4C). This demonstrates that s5PSeq combined with nanopore sequencing can enable phenotypic molecular testing for AMR in clinical labs. In addition to antimicrobial sensitivity testing, being a sequencing-based approach, in principle s5PSeq can also provide information on strain identification and presence of AMR genes.

## Discussion

In this study, we have developed s5PSeq, a method that simplifies the study of 5’P mRNA degradome by reducing costs, manual effort, and library preparation time. We have demonstrated a novel approach for rRNA depletion that selectively inhibits the single-stranded RNA ligation of 5’P rRNA molecules, and eliminates the necessity for expensive and time-consuming rRNA depletion procedures. Lastly, we show that s5PSeq can be easily integrated with nanopore sequencing reducing turnaround time.

To illustrate s5PSeq applicability, we study its ability to capture changes in ribosome dynamics after antibiotic treatment^9,10^. We focused on ribosome-targeting antibiotics in bacteria with RNase J, where 5PSeq provides single-nucleotide resolution information and ribosome stalls can be easily connected to the drug’s mechanism of action^9^. We demonstrate that 5’P mRNA degradation signatures can serve as a rapid molecular readout for antimicrobial susceptibility. By quantifying context-specific ribosome stalls^9,10^ after short (10 minutes) antibiotic exposure, s5PSeq provides a functional readout of translation inhibition that correlates with long-term bacterial growth. Our findings show that these drug- and context-specific ribosome stalling RNA degradation signatures can differentiate the effects of various antibiotic concentrations (Figure 1) and distinguish *Clostridioides difficile* resistant strains from sensitive ones (Figures 3 and 4). We show that combining s5PSeq with nanopore sequencing further enhances its clinical utility by enabling real-time data acquisition and reducing turnaround time to within a single working shift (RNA to Fastq). s5PSeq can be performed using the cost-effective Flongle flow cells and MinION devices, which are already available in many clinical microbiology laboratories. Our current implementation enables rapid, cost-effective, and accurate profiling of phenotypic AMR, with results delivered within 10 hours (starting from metabolically active bacteria). Additionally, even using our current implementation this turnaround time can be further reduced by utilizing flow cells with higher throughput per unit time. For instance, instead of multiplexing 24 samples per Flongle and running sequencing for 5 hours, theoretically we could obtain sufficient coverage for sample discrimination (e.g., minimum of 3,000 mapped reads as in Figure 4d) by multiplexing 32 samples in a conventional MinION flow cell for 1hour. This would reduce the sample-to-answer time to less than 6 hours, fitting within a typical clinical working shift. In fact, during the preparation of this manuscript ONT has decided to discontinue Flongle flow cells in favor of regular MinION flow cells, which offer higher throughput and sequencing quality. Further optimization of the s5PSeq experimental and computational protocol is expected to reduce the time required.

Detecting ribosomal stalling patterns induced by ribosome-targeting antibiotics provides a phenotypic readout of the drug activity and reflects the actual impact of antibiotic treatment. This approach bridges the gap between traditional phenotypic assays and molecular diagnostic approaches. Furthermore, being a sequencing-based approach, s5PSeq does not only provide phenotypic antimicrobial sensitivity information but also genetic information, enabling the identification of specific strains or the presence of AMR marker genes. Unlike other phenotypic approaches that measure cell growth at the cellular level (e.g., traditional microbial cultures or microscopy-based approaches)^30^, in principle 5’P mRNA degradome analysis does not require strain isolation and can be applied to complex samples^9^. This feature has the potential to further reduce the time from sample collection to antimicrobial sensitivity testing in real-life scenarios by reducing or even eliminating the time for isolate identification. However, further research is necessary to assess s5PSeq applicability to real-life complex samples within the clinical setting, such as patient-derived blood cultures. The potential of s5PSeq extends beyond *C.difficile*, and can be easily extended to other pathogens with 5’-3’ co-translational mRNA decay^9,31^. In *C.difficile* erythromycin resistance is widespread^32^, and phenotypic testing for this antibiotic is performed mainly for antimicrobial resistance surveillance and susceptibility trend prediction. Thus, future research should focus on expanding the approach to a wider range of pathogens and antibiotics (e.g., *E.faecium*, *S.aureus*…).

In conclusion, we believe that the integration of s5PSeq method with nanopore sequencing paves the way for the development of rapid, cost-effective, and accurate approach to phenotypic antimicrobial resistance diagnostics. This innovative strategy effectively addresses critical limitations in current diagnostic methods and holds significant potential to enhance clinical decision-making, improve patient care, and contribute to global public health efforts against antimicrobial resistance.

### Limitations of the study

While s5PSeq offers a rapid and cost-effective approach for studying the 5’P degradome and profiling phenotypic antimicrobial resistance, several limitations should be considered. First, although 5’P sequencing provides single-nucleotide resolution for species possessing 5’-3’ exonucleases (e.g., RNase J), it remains essential to perform preliminary studies characterizing antibiotic-specific ribosome stalls across varying times and drug concentrations, and to correlate these signatures with differential bacterial growth. While s5PSeq could theoretically be employed to examine general translational perturbations in response to non-ribosome-targeting drugs^31^ or to study species lacking a 5’-3’ RNA exonuclease, where s5PSeq resolution is lower, further optimization is necessary.

Second, while s5PSeq has the potential to be applied directly to complex clinical samples without prior strain isolation, our current study was limited to pure cultures. Even if sensitivity of s5PSeq relies primarily on sequencing depth, additional work will be required to demonstrate its ability to identify alterations in ribosome dynamics in mixed or low-abundance bacterial samples.

Third, s5PSeq can theoretically detect drug-induced changes in any metabolically active bacteria. However, further research is required to demonstrate the utility of s5PSeq in real-world clinical samples such as patient-derived blood cultures prior to strain isolation.

Finally, successful implementation of s5PSeq in clinical settings will require the development of robust sample-to-answer analysis pipelines, both experimental and computational, that meet strict *in vitro* diagnostic regulations. Comprehensive benchmarking against existing antimicrobial susceptibility tests in clinical settings will also be critical.

## Supporting information

Supplementary protocol

Interactive s5PSeq

## Resource availability

### Lead contact

Requests for further information or resources should be directed to the lead contact, Vicent Pelechano

### Materials availability

No new unique reagents have been generated in this study.

### Data and code availability

Sequencing data will been deposited on GEO. Table S3 includes a list of all samples sequenced.

## Acknowledgements

We wish to thank members of the Pelechano, Du, Kutter and Friedländer laboratories for useful discussions. This project was mainly funded by a Research collaboration grant China-Sweden from the National Natural Science Foundation of China [82161138017] and the Swedish Research Council [VR 2021-06112] to VP, JD and WHC. Additional funding is acknowledge<u>d</u> from the Swedish Research Council [VR 2022-05272, 2023-02026 and 2024-03210], an extension Wallenberg Academy Fellowship [2021.0167] and Karolinska Institutet (SciLifeLab, SFO, KID and KI funds) to VP.; the Swedish Research Council [2021-01683 and 2021-06112] and Cancerfonden [23 2916 Pj] to JD.; the EU H2020-MSCA-IF-2018 program under grant agreement [845495 - TERMINATOR] and the MoESCS RA HESC grant [24FP-2I061] to LN; Computational analysis was enabled by resources provided by the National Academic Infrastructure for Supercomputing in Sweden (NAISS), partially funded by the Swedish Research Council through grant agreement no. 2022-06725. We acknowledge support from the National Genomics Infrastructure in Stockholm funded by Science for Life Laboratory, the Knut and Alice Wallenberg Foundation and the Swedish Research Council for the PromethION Sequencing, and NAISS for assistance with massively parallel sequencing and access to the UPPMAX and PDC computational infrastructure.

## Author contributions

VP, HL and SH conceived and designed the study. HL performed the bulk of the experimental work with support from RH, FRG and SH. HL performed data analysis with support from RH and LN. XY, WHC and JD contributed to data interpretation, design and supervision. HL, and VP drafted the initial manuscript and all authors revised it. VP supervised the study.

## STAR★Methods

### Key resources table

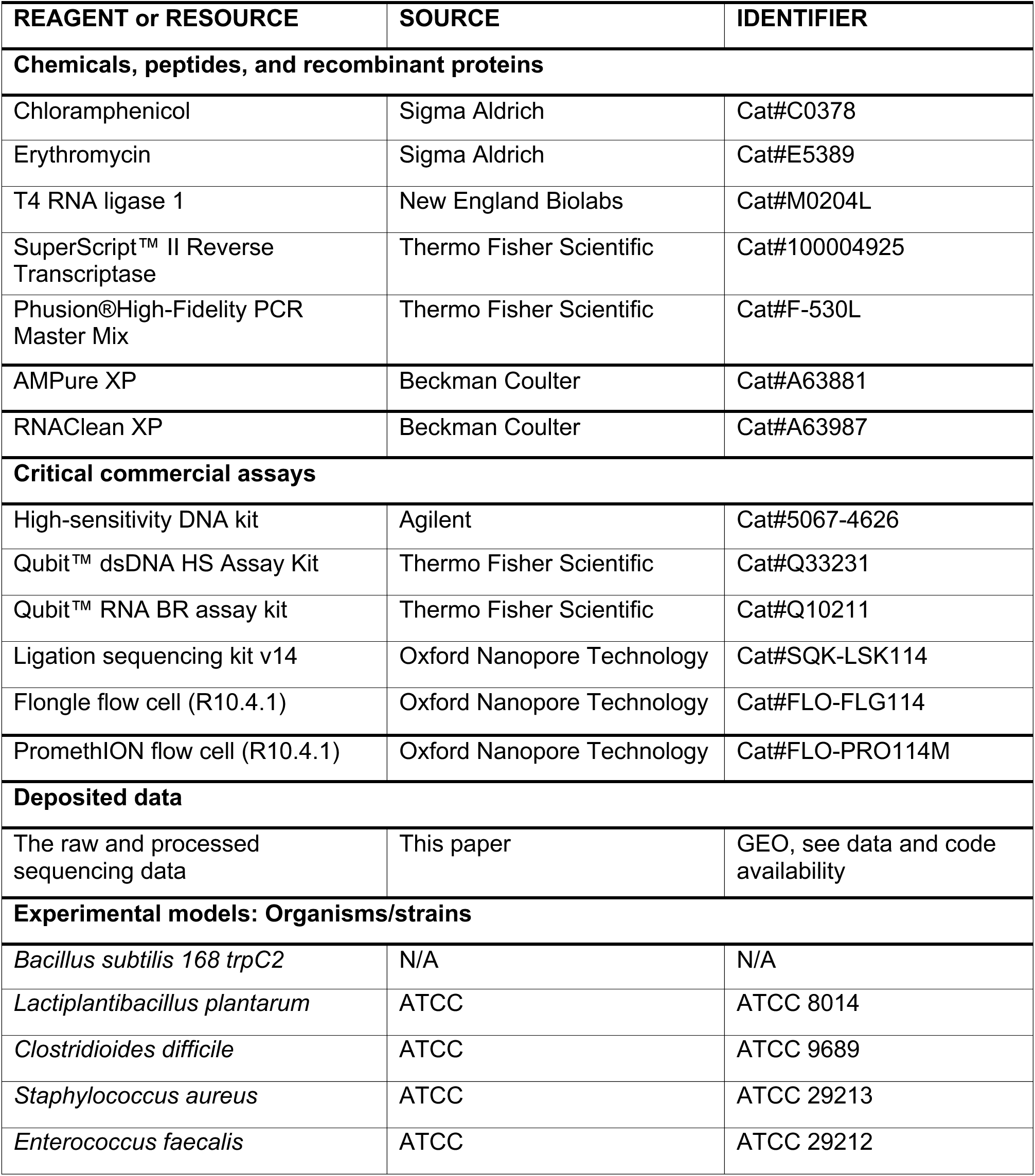
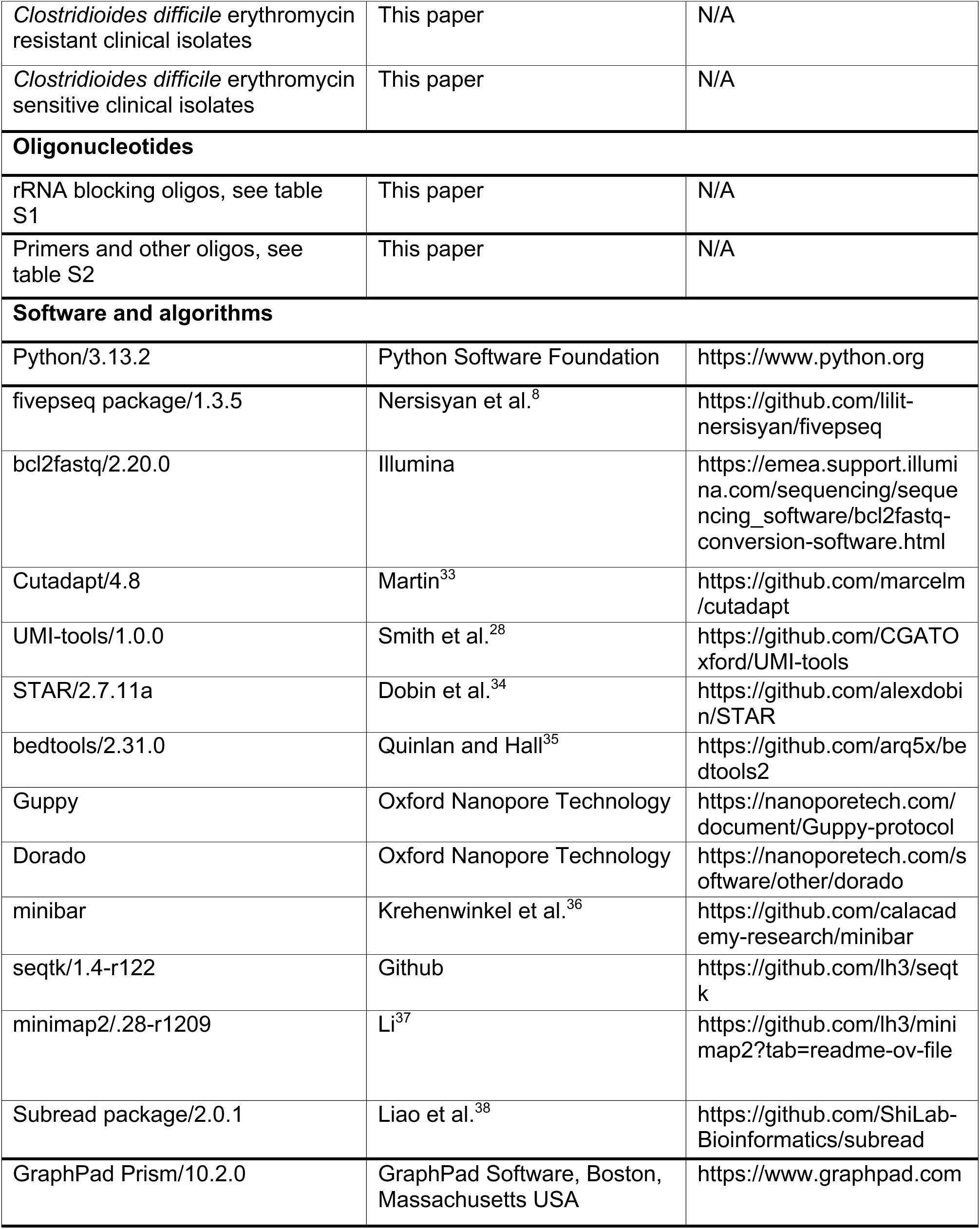

### Experimental model and study participant details

#### Bacterial Strains and Culture Conditions

*Bacillus subtilis* 168 trpC2 and *Staphylococcus aureus* ATCC 29213 were cultured aerobically at 37 °C in LB medium (Sigma-Aldrich, Cat# L7275). *Lactiplantibacillus plantarum* ATCC 8014 was cultured aerobically at 37 °C in MRS medium (Sigma-Aldrich, Cat#69966). *Enterococcus faecalis* ATCC 29212 was grown anaerobically at 37 °C in NYCIII medium. *Clostridioides difficile* ATCC 9689, along with erythromycin-resistant and -sensitive *C. difficile* clinical isolates, were cultured anaerobically at 37 °C in brain heart infusion (BHI) medium<Edwards, 2013 #44>. Two C. difficile clinical isolates (one erythromycin-resistant and one erythromycin-sensitive) were obtained from the Public Health Agency of Sweden (Dnr 03016-2017 1.4.4 and 03016-2022). These isolates were cultured under anaerobic conditions at 37 °C in BHI medium.

#### Antibiotic Treatment and Growth Assays

To evaluate antibiotic sensitivity via standard growth assays, overnight cultures of *Bacillus subtilis* and *Lactiplantibacillus plantarum* were diluted to an initial OD₆₀₀ of 0.005–0.05 and incubated aerobically in 96-well plates for 18–24 hours in the presence of chloramphenicol (0, 1.6, 8, or 40 µg/mL) or erythromycin (0, 0.1, 0.5, or 2.5 µg/mL), with growth monitored using a Tecan Spark® microplate reader. For *Clostridioides difficile* clinical isolates, overnight cultures were diluted to an OD₆₀₀ of 0.05 and incubated anaerobically in BHI medium with erythromycin (0, 1, or 256 µg/mL) under the same conditions. Growth was measured using a Tecan Sunrise™ microplate reader.

#### Preparation of RNA Samples for HT-5PSeq and s5PSeq

For preparation of metabolically active cultures used in HT-5PSeq or s5PSeq library construction, overnight cultures of *Bacillus subtilis* and *Lactiplantibacillus plantarum* were diluted and grown to mid-exponential phase (OD₆₀₀ = 0.6–0.8) over 4–8 hours. A 1 mL aliquot was treated with the same concentrations of chloramphenicol or erythromycin as used in the growth assays for 10 minutes, followed by centrifugation at 13,000 rpm for 3 minutes. The supernatant was discarded, and the resulting cell pellets were frozen on dry ice for RNA extraction. Exponentially growing cultures of *Enterococcus faecalis* ATCC 29212, *Staphylococcus aureus* ATCC 29213, and *Clostridioides difficile* ATCC 9689 were processed using the same procedure to evaluate the s5PSeq protocol across multiple bacterial species. Similarly, overnight cultures of *C. difficile* clinical isolates were diluted to an OD₆₀₀ of 0.05 and grown to mid-exponential phase (OD₆₀₀ = 0.6–0.8) over 7–8 hours. A 1 mL aliquot was treated with erythromycin (0 or 1 µg/mL) for 10 minutes, followed by the addition of 500 µL RNAprotect Bacteria Reagent (QIAGEN). After a 5-minute incubation, cells were harvested by centrifugation at 13,000 rpm for 3 minutes, and pellets were frozen on dry ice for RNA extraction.

### Method details

#### RNA extraction

Total RNA was extracted using a previously established phenol-chloroform method combined with beads-beating^9^. Briefly, frozen cell pellets were resuspended in a lysis reagent with phenol and glass beads, followed by mechanical disruption using a MultiMixer for 2 minutes or a FastPrep^®^ system for 1min. The aqueous phase was then separated and further purified using acidic phenol-chloroform extraction. Finally, total RNA was precipitated in ethanol and eluted in nuclease-free water. RNA concentrations were quantified using a Nanodrop spectrophotometer and Qubit fluorometer. RNA quality was assessed by gel electrophoresis and/or an Agilent Bioanalyzer using the RNA Pico kit. The extracted RNA showed high integrity with minimal DNA contamination.

#### s5PSeq Library Preparation

s5PSeq libraries were prepared from 200 ng of total RNA without DNase treatment. Ribosomal RNA was blocked by hybridization with 0.4 µM of a mixture of 146 universal blocking oligonucleotides and 4 µM of two species-specific oligonucleotides targeting the 5′ ends of 16S and 23S rRNAs (Table S1), in the presence of 50 mM NaCl. The hybridization reaction was performed using the following thermal profile: 75 °C for 5 min, 65 °C for 5 min, followed by a stepwise temperature decrease of 3 °C every 2 min until 35 °C. After rRNA blocking, 1 µM of single-stranded RNA adapters containing unique molecular identifiers (UMIs) and barcodes were ligated to RNA using T4 RNA ligase 1 in the presence of 1 mM ATP and 1X T4 RNA ligation buffer at 25 °C for 1 hour. Ligated RNA was purified using RNA CleanXP beads (1.8X volume). First-strand cDNA synthesis was then performed using SuperScript II reverse transcriptase and random hexamers following the manufacturer’s protocol. Template RNA was degraded by incubation with 28.6 mM NaOH at 65 °C for 20 min, followed by neutralization with an equimolar amount of Tris-HCl. cDNA was purified using Ampure XP beads (1.8X volume). Libraries were amplified using 0.25 µM Illumina-compatible primers with dual barcodes and 2X Phusion High-Fidelity PCR Master Mix with the following cycling conditions: 98 °C for 30 s; 15 cycles of 98 °C for 20 s, 65 °C for 30 s, and 72 °C for 30 s; followed by 72 °C for 7 min. PCR products were size-selected using a two-step selection with Ampure XP beads (0.6X–0.8X) to enrich for fragments between 400–600 bp. All oligonucleotide sequences used in the protocol are listed in Table S2.

#### Illumina Sequencing and Data Analysis

Size-selected libraries were quantified using Qubit dsDNA HS assays, and 650 pM libraries were loaded onto the Illumina NextSeq 2000 platform for sequencing. Demultiplexing of Illumina reads for each sample was performed using bcl2fastq. Preprocessing, including adapter trimming and quality filtering with cutadapt^33^, UMI extraction with UMI-tools^28^ were conducted using previously established workflows^8^. Reads were aligned to the respective bacterial reference genomes using STAR^34^, and mapped reads were analyzed using the fivepseq package^8^ to identify ribosomal stalling patterns and phenotypic antimicrobial resistance (AMR) signatures.

#### Nanopore Sequencing and Data Analysis

Libraries originally prepared for Illumina sequencing were repurposed for nanopore sequencing by adding nanopore adapters using the Oxford Nanopore Technologies ligation sequencing kit v14 (SQK-LSK114). Libraries were loaded onto a Flongle flow cell and sequenced on the MinION platform. The same libraries were also loaded onto a PromethION flow cell and sequenced on the PromethION 48 platform. Raw nanopore sequencing data generated in pod5 format were retrieved after 24 hours of sequencing on a MinION flongle flow cell or 72 hours of sequencing on a PromethION flow cell. Base calling was performed using Dorado software in Fast base calling model for flongle sequencing, or real-time base calling with Guppy software in Super accurate (SUP) base calling model. Reads were demultiplexed based on the dual barcode Illumina adapter using the minibar.py program^39^. UMI extraction was performed using UMI-tools^28^, followed by alignment to the *C. difficile* reference genome using minimap2^37^. Mapped reads were analyzed using the fivepseq package^8^ to identify ribosomal stalling patterns and phenotypic AMR signatures.

### Quantification and statistical analysis

5′P degradome data were analyzed using the fivepseq package (v1.3.5)^8^ to generate codon-resolution coverage profiles and metagene plots. Ribosome stalling was quantified as the ratio of 5′P signal intensity at defined positions upstream of specific codons to the average signal in the surrounding regions, with signals normalized as reads per million (RPM) of mapped reads. Statistical comparisons were performed using unpaired two-tailed *t*-tests (*p* < 0.0001), and reproducibility was assessed by Pearson correlation. Data visualizations were generated using GraphPad Prism 10 for amino acid-specific stalling plots and heatmap, metagene coverage at start and stop codons, and RNA composition bar plots based on fivepseq output.

### Declaration of generative AI and AI-assisted technologies in the writing process

During the preparation of this work the authors used Microsoft Copilot in order to proofread the text. After using this tool, the authors reviewed and edited the content as needed and take full responsibility for the content of the publication.

## Reference

1. Naddaf, M. (2024). 40 million deaths by 2050: toll of drug-resistant infections to rise by 70. Nature 633, 747–748. 10.1038/d41586-024-03033-w.

2. Clsi. (2018). Methods for dilution antimicrobial susceptibility tests for bacteria that grow aerobically. 10 th ed. CLSI standard M07.

3. Redding, L.E., Tu, V., Abbas, A., Alvarez, M., Zackular, J.P., Gu, C., Bushman, F.D., Kelly, D.J., Barnhart, D., Lee, J.J., and Bittinger, K.L. (2022). Genetic and phenotypic characteristics of Clostridium (Clostridioides) difficile from canine, bovine, and pediatric populations. Anaerobe 74, 102539. 10.1016/j.anaerobe.2022.102539.

4. Anjum, M.F., Zankari, E., and Hasman, H. (2017). Molecular Methods for Detection of Antimicrobial Resistance. Microbiol Spectr 5. 10.1128/microbiolspec.ARBA-0011-2017.

5. Rebelo, A.R., Bortolaia, V., Leekitcharoenphon, P., Hansen, D.S., Nielsen, H.L., Ellermann-Eriksen, S., Kemp, M., Røder, B.L., Frimodt-Møller, N., Søndergaard, T.S., et al. (2022). One Day in Denmark: Comparison of Phenotypic and Genotypic Antimicrobial Susceptibility Testing in Bacterial Isolates From Clinical Settings. Frontiers in Microbiology Volume 13 - 2022. 10.3389/fmicb.2022.804627.

6. Pelechano, V., Wei, W., and Steinmetz, L.M. (2015). Widespread Co-translational RNA Decay Reveals Ribosome Dynamics. Cell 161, 1400–1412. 10.1016/j.cell.2015.05.008.

7. Zhang, Y., and Pelechano, V. (2021). High-throughput 5’P sequencing enables the study of degradation-associated ribosome stalls. Cell Rep Methods 1, 100001. 10.1016/j.crmeth.2021.100001.

8. Nersisyan, L., Ropat, M., and Pelechano, V. (2020). Improved computational analysis of ribosome dynamics from 5’P degradome data using fivepseq. NAR Genom Bioinform 2, lqaa099. 10.1093/nargab/lqaa099.

9. Huch, S., Nersisyan, L., Ropat, M., Barrett, D., Wu, M., Wang, J., Valeriano, V.D., Vardazaryan, N., Huerta-Cepas, J., Wei, W., et al. (2023). Atlas of mRNA translation and decay for bacteria. Nat Microbiol 8, 1123–1136. 10.1038/s41564-023-01393-z.

10. Crowe-McAuliffe, C., Murina, V., Turnbull, K.J., Huch, S., Kasari, M., Takada, H., Nersisyan, L., Sundsfjord, A., Hegstad, K., Atkinson, G.C., et al. (2022). Structural basis for PoxtA-mediated resistance to phenicol and oxazolidinone antibiotics. Nat Commun 13, 1860. 10.1038/s41467-022-29274-9.

11. Zhang, Y., and Pelechano, V. (2021). Application of high-throughput 5’P sequencing for the study of co-translational mRNA decay. STAR Protoc 2, 100447. 10.1016/j.xpro.2021.100447.

12. Balouiri, M., Sadiki, M., and Ibnsouda, S.K. (2016). Methods for in vitro evaluating antimicrobial activity: A review. J Pharm Anal 6, 71–79. 10.1016/j.jpha.2015.11.005.

13. Ismail, B.B., Wang, W., Ayub, K.A., Guo, M., and Liu, D. (2024). Advances in microscopy-based techniques applied to the antimicrobial resistance of foodborne pathogens. Trends in Food Science & Technology 152, 104674. 10.1016/j.tifs.2024.104674.

14. Spears, J.L., Kramer, R., Nikiforov, A.I., Rihner, M.O., and Lambert, E.A. (2021). Safety Assessment of Bacillus subtilis MB40 for Use in Foods and Dietary Supplements. Nutrients 13. 10.3390/nu13030733.

15. Larsen, N., Thorsen, L., Kpikpi, E.N., Stuer-Lauridsen, B., Cantor, M.D., Nielsen, B., Brockmann, E., Derkx, P.M., and Jespersen, L. (2014). Characterization of Bacillus spp. strains for use as probiotic additives in pig feed. Appl Microbiol Biotechnol 98, 1105–1118. 10.1007/s00253-013-5343-6.

16. Rojo-Bezares, B., Saenz, Y., Poeta, P., Zarazaga, M., Ruiz-Larrea, F., and Torres, C. (2006). Assessment of antibiotic susceptibility within lactic acid bacteria strains isolated from wine. Int J Food Microbiol 111, 234–240. 10.1016/j.ijfoodmicro.2006.06.007.

17. Marks, J., Kannan, K., Roncase, E.J., Klepacki, D., Kefi, A., Orelle, C., Vazquez-Laslop, N., and Mankin, A.S. (2016). Context-specific inhibition of translation by ribosomal antibiotics targeting the peptidyl transferase center. Proc Natl Acad Sci U S A 113, 12150–12155. 10.1073/pnas.1613055113.

18. Davis, A.R., Gohara, D.W., and Yap, M.N. (2014). Sequence selectivity of macrolide-induced translational attenuation. Proc Natl Acad Sci U S A 111, 15379–15384. 10.1073/pnas.1410356111.

19. Krawczyk, S.J., Lesniczak-Staszak, M., Gowin, E., and Szaflarski, W. (2024). Mechanistic Insights into Clinically Relevant Ribosome-Targeting Antibiotics. Biomolecules 14. 10.3390/biom14101263.

20. Sothiselvam, S., Neuner, S., Rigger, L., Klepacki, D., Micura, R., Vazquez-Laslop, N., and Mankin, A.S. (2016). Binding of Macrolide Antibiotics Leads to Ribosomal Selection against Specific Substrates Based on Their Charge and Size. Cell Rep 16, 1789–1799. 10.1016/j.celrep.2016.07.018.

21. Kannan, K., Kanabar, P., Schryer, D., Florin, T., Oh, E., Bahroos, N., Tenson, T., Weissman, J.S., and Mankin, A.S. (2014). The general mode of translation inhibition by macrolide antibiotics. Proc Natl Acad Sci U S A 111, 15958–15963. 10.1073/pnas.1417334111.

22. Alekseenko, A., Barrett, D., Pareja-Sanchez, Y., Howard, R.J., Strandback, E., Ampah-Korsah, H., Rovsnik, U., Zuniga-Veliz, S., Klenov, A., Malloo, J., et al. (2021). Direct detection of SARS-CoV-2 using non-commercial RT-LAMP reagents on heat-inactivated samples. Sci Rep 11, 1820. 10.1038/s41598-020-80352-8.

23. Pelechano, V., Wei, W., and Steinmetz, L.M. (2016). Genome-wide quantification of 5’-phosphorylated mRNA degradation intermediates for analysis of ribosome dynamics. Nat Protoc 11, 359–376. 10.1038/nprot.2016.026.

24. Bartlett, J.G. (1994). Clostridium difficile: history of its role as an enteric pathogen and the current state of knowledge about the organism. Clin Infect Dis 18 *Suppl 4*, S265–272. 10.1093/clinids/18.supplement_4.s265.

25. Tilkorn, F., Frickmann, H., Simon, I.S., Schwanbeck, J., Horn, S., Zimmermann, O., Gross, U., Bohne, W., and Zautner, A.E. (2020). Antimicrobial Resistance Patterns in Clostridioides difficile Strains Isolated from Neonates in Germany. Antibiotics (Basel) 9. 10.3390/antibiotics9080481.

26. Peng, Z., Jin, D., Kim, H.B., Stratton, C.W., Wu, B., Tang, Y.W., and Sun, X. (2017). Update on Antimicrobial Resistance in Clostridium difficile: Resistance Mechanisms and Antimicrobial Susceptibility Testing. J Clin Microbiol 55, 1998–2008. 10.1128/JCM.02250-16.

27. Freeman, J., Sanders, I., Harmanus, C., Clark, E.V., Berry, A.M., and Smits, W.K. (2025). Antimicrobial susceptibility testing of Clostridioides difficile: a dual-site study of three different media and three therapeutic antimicrobials. Clin Microbiol Infect. 10.1016/j.cmi.2025.01.028.

28. Smith, T., Heger, A., and Sudbery, I. (2017). UMI-tools: modeling sequencing errors in Unique Molecular Identifiers to improve quantification accuracy. Genome Res 27, 491–499. 10.1101/gr.209601.116.

29. Payne, A., Holmes, N., Clarke, T., Munro, R., Debebe, B.J., and Loose, M. (2021). Readfish enables targeted nanopore sequencing of gigabase-sized genomes. Nat Biotechnol 39, 442–450. 10.1038/s41587-020-00746-x.

30. Jorgensen, J.H., and Ferraro, M.J. (2009). Antimicrobial susceptibility testing: a review of general principles and contemporary practices. Clin Infect Dis 49, 1749–1755. 10.1086/647952.

31. Stevens, I., Silao, F.G., Huch, S., Liu, H., Ryman, K., Carvajal-Jimenez, A., Ljungdahl, P.O., and Pelechano, V. (2024). The early transcriptional and post-transcriptional responses to fluconazole in sensitive and resistant Candida albicans. Sci Rep 14, 29012. 10.1038/s41598-024-80435-w.

32. Dilnessa, T., Getaneh, A., Hailu, W., Moges, F., and Gelaw, B. (2022). Prevalence and antimicrobial resistance pattern of Clostridium difficile among hospitalized diarrheal patients: A systematic review and meta-analysis. PLoS One 17, e0262597. 10.1371/journal.pone.0262597.

33. Martin, M. (2011). Cutadapt removes adapter sequences from high-throughput sequencing reads. 2011 *17*, 3. 10.14806/ej.17.1.200.

34. Dobin, A., Davis, C.A., Schlesinger, F., Drenkow, J., Zaleski, C., Jha, S., Batut, P., Chaisson, M., and Gingeras, T.R. (2013). STAR: ultrafast universal RNA-seq aligner. Bioinformatics 29, 15–21. 10.1093/bioinformatics/bts635.

35. Quinlan, A.R., and Hall, I.M. (2010). BEDTools: a flexible suite of utilities for comparing genomic features. Bioinformatics 26, 841–842. 10.1093/bioinformatics/btq033.

36. Krehenwinkel, H., Pomerantz, A., Henderson, J.B., Kennedy, S.R., Lim, J.Y., Swamy, V., Shoobridge, J.D., Graham, N., Patel, N.H., Gillespie, R.G., and Prost, S. (2019). Nanopore sequencing of long ribosomal DNA amplicons enables portable and simple biodiversity assessments with high phylogenetic resolution across broad taxonomic scale. GigaScience 8, giz006. 10.1093/gigascience/giz006.

37. Li, H. (2018). Minimap2: pairwise alignment for nucleotide sequences. Bioinformatics 34, 3094–3100. 10.1093/bioinformatics/bty191.

38. Liao, Y., Smyth, G.K., and Shi, W. (2013). The Subread aligner: fast, accurate and scalable read mapping by seed-and-vote. Nucleic Acids Res 41, e108. 10.1093/nar/gkt214.

39. Krehenwinkel, H., Pomerantz, A., Henderson, J.B., Kennedy, S.R., Lim, J.Y., Swamy, V., Shoobridge, J.D., Graham, N., Patel, N.H., Gillespie, R.G., and Prost, S. (2019). Nanopore sequencing of long ribosomal DNA amplicons enables portable and simple biodiversity assessments with high phylogenetic resolution across broad taxonomic scale. Gigascience 8. 10.1093/gigascience/giz006.

