## Supplementary files for "A streamlined nanopore-compatible 5PSeq protocol for rapid phenotypic antimicrobial sensitivity testing"

### Supplementary material

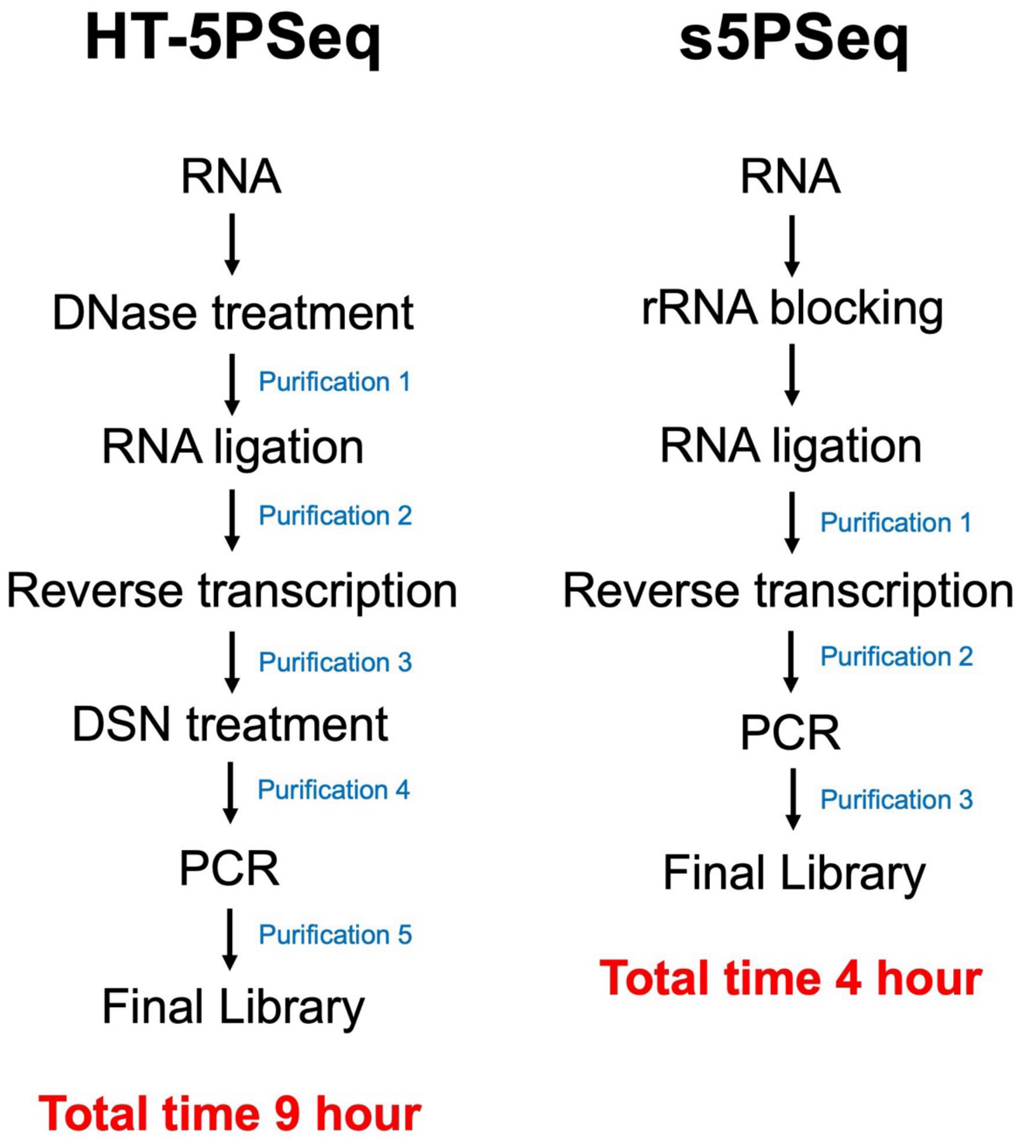


**Figure S1. Comparison between HT-5PSeq and s5PSeq protocol.** The simplified s5PSeq method streamlines library preparation by reducing pipetting steps and minimizing purification, shortening the total workflow from 9 hours to 4 hours.

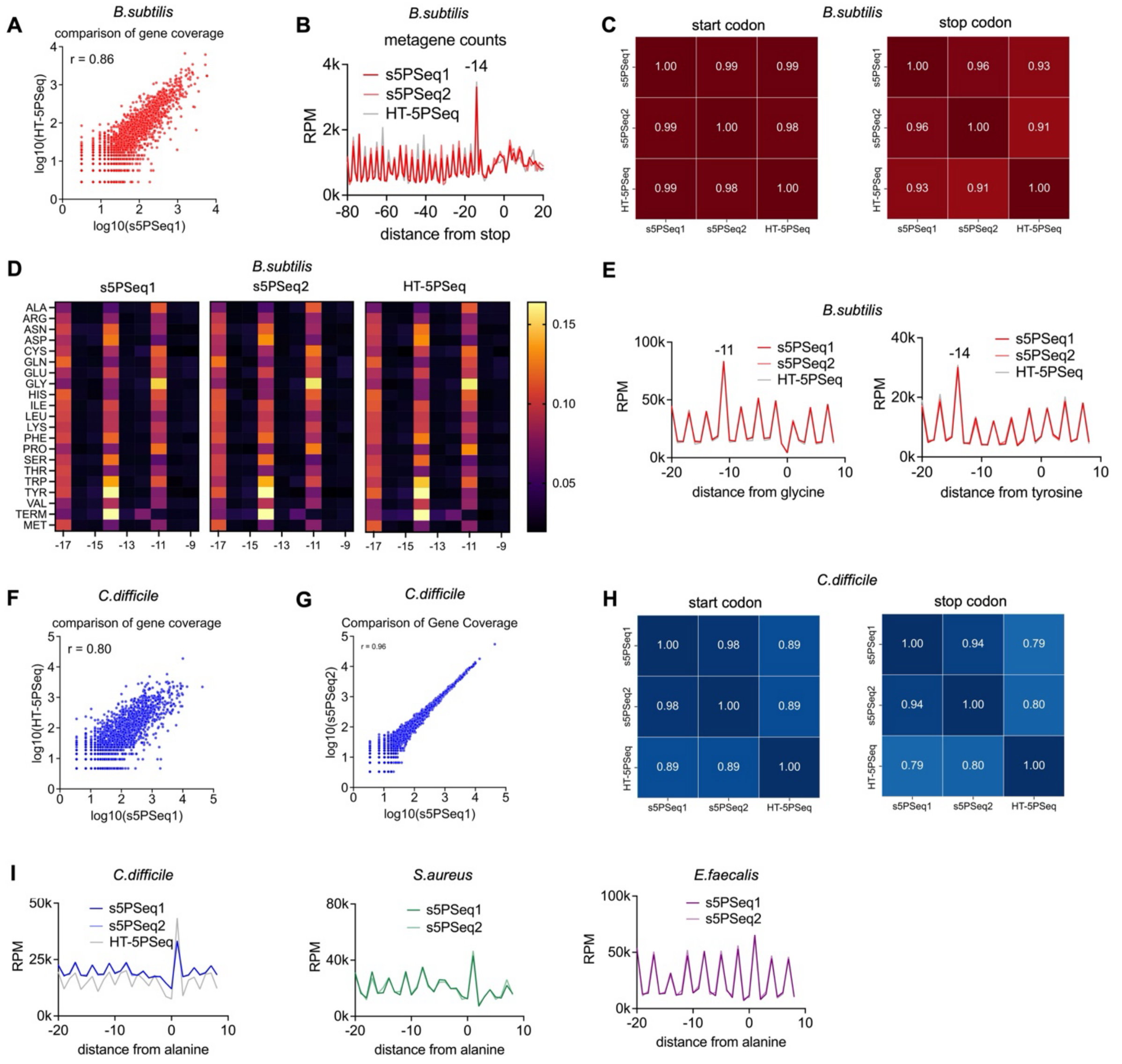


**Figure S2. Additional quality controls for s5Pseq method**

(A) Gene coverage comparison between s5PSeq1 and HT-5PSeq in *B.subtilis*, showing a strong linear correlation (Pearson's *r* = 0.86). Gene counts were normalized using median ratio normalization, followed by $\log_{10}$ transformation for visualization.

(B) Metagene analysis of 5'P mRNA read coverage relative to the stop codons in *B. subtilis*.

(C) Heatmap of Pearson correlation for normalized 5′P metagene coverage (RPM) around start codon (−50 to +50 bp) and stop codons (−80 to +20 bp) in *B. subtilis,* showing strong agreement between HT-5PSeq and s5PSeq replicates (s5PSeq1 and s5PSeq2).

(D) Heatmap showing amino acid-specific 5′P coverage. Positions at 14 and 11 nucleotides upstream of the amino acid codons correspond to ribosome occupancy at the A and P sites, respectively. For each amino acid codons, reads were normalized to the total number of 5′P reads within the −20 to −1 nucleotide window.

(E) Line plot showing 5’P mRNA degradation profiles at amino acid codon specific positions (x axis showing relative distance to glycine and tyrosine codons respectively) obtained using s5Pseq (s5PSeq1 and s5PSeq2) and HT-5Pseq in *B. subtilis.*

(F) Gene coverage comparison between s5PSeq1 and HT-5PSeq in *C.difficile* as in (A), showing a strong linear correlation (Pearson’s $r$ = 0.80). Gene counts were normalized using median ratio normalization, followed by $\log_{10}$ transformation for visualization.

(G) Gene coverage comparison between replicates s5PSeq1 and s5PSeq2 in *C.difficile*, showing a strong linear correlation (Pearson’s $r$ = 0.96). Gene counts were normalized using median ratio normalization, followed by $\log_{10}$ transformation for visualization.

(H) Heatmap of Pearson correlation for normalized 5′P metagene coverage (RPM) around start codon (−50 to +50 bp) and stop codons (−80 to +20 bp) in *C.difficile* as in (C), showing strong agreement between HT-5PSeq and s5PSeq replicates (s5PSeq1 and s5PSeq2).

(I) Line plot showing 5’P mRNA degradation profiles at amino acid codon specific positions (x axis showing relative distance to alanine codons) in *C.difficile*, *S.aureus* and *E.faecalis,* as in (E).

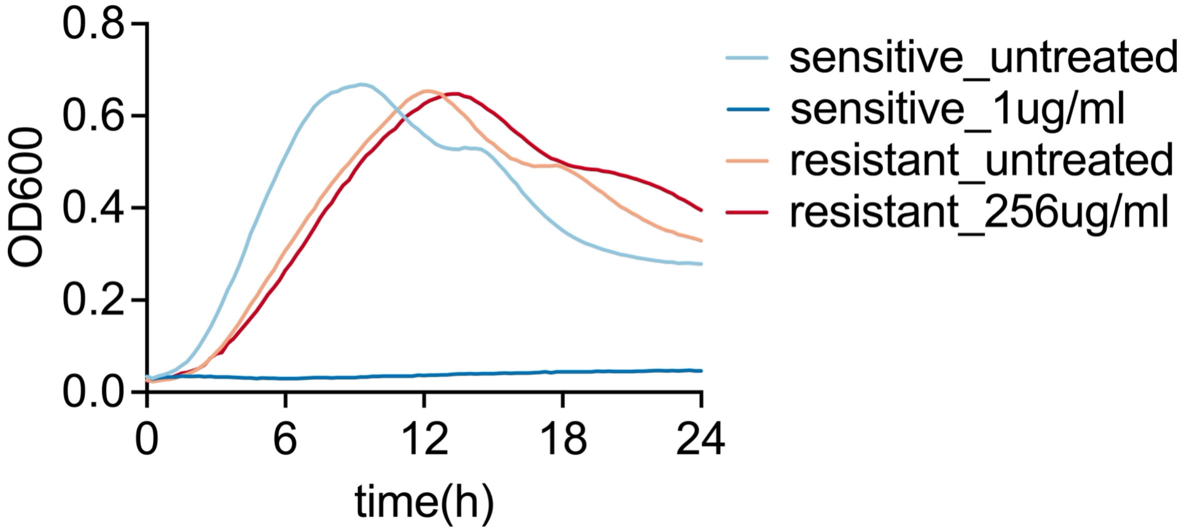


**Figure S3. Growth curve of Clostridium difficile clinical isolates.** Erythromycin-resistant and –sensitive strains were grown in presence (treated) and absence (untreated) erythromycin at concentrations of 1 μg/mL and 256 μg/mL. Growth of the erythromycin-sensitive strain is completely suppressed at 1 μg/mL, whereas the resistant strain exhibits robust growth at 256 μg/mL erythromycin concentration.

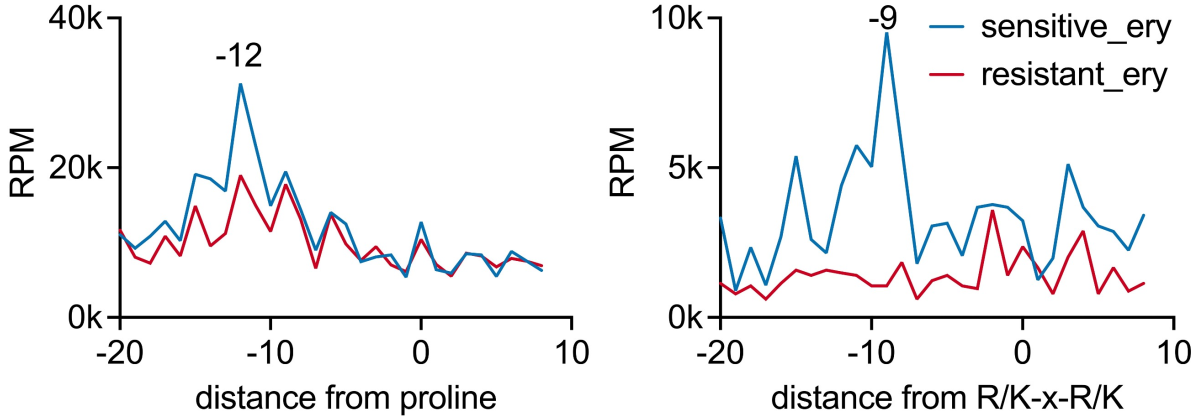


**Figure S4. PromethION subsampling for s5PSeq**. Line plots showing erythromycin-induced ribosomal stalling, with clear differentiation between erythromycin-sensitive and -resistant *C.difficile* strains when 30,000 reads are subsampled from Fig 4a. Stalling occurs at positions 12 nucleotides upstream of proline residues and 9 nucleotides upstream of the R/K-x-R/K motif (with the first base of the final codon in the motif set at position 0).

**Table S1.** s5PSeq rRNA blocking oligonucleotides.

**Table S2.** Other used oligonucleotides.

**Table S3.** Summary of sequenced samples.

**Suppelemntary  file.** Interactive FivePSeq report

**Supplementary protocol.** Detailed s5PSeq protocol
