## Supplementary protocol for "A streamlined nanopore-compatible 5PSeq protocol for rapid phenotypic antimicrobial sensitivity testing"

### Protocol: Simplified 5PSeq(s5PSeq) Library Preparation

**Starting Material:** 200ng total bacterial RNA

#### 1. rRNA Blocking Oligo Hybridization (30min)

In a 200 µL PCR tube, add the following reagents. Mix by gently flicking the tube and briefly spin down. Run the following program on a thermal cycler: 75 °C for 5 min; 65 °C for 5 min, and stepwise decrease of 3 °C every 2 min to 35 °C.

| Reagent | Volume(µL) |
|---|---|
| Bacterial RNA (200ng) | 1 |
| Blocking oligo_16s5p (20µM) * | 1 |
| Blocking oligo_23s5p (20µM) * | 1 |
| Blocking oligo_universal (2µM) ** | 1 |
| NaCl solution(250mM) | 1 |
| Total | 5 |

*Species-specific oligos (see Table S2).
**Mixture of 146 oligos (see Table S2).

#### 2. ssRNA Ligation (1hour)

To the hybridized sample, add the following reagents. Mix by flicking the tube and briefly spin down. Incubate at 25 °C for 1 hour with the lid temperature off.

| Reagent | Volume(µL) |
|---|---|
| rP5_RND adapter (10µM) | 1 |
| 10X T4 RNA ligation buffer | 1 |
| T4 RNA ligase 1 (10U/µL) *** | 1 |
| Nuclease-free H2O | 1 |
| ATP (10mM) | 1 |
| Sample from Step 1 | 5 |
| Total | 10 |

*** T4 RNA ligase 1 can be substituted with other single-stranded RNA ligases with comparable 5′-phosphate RNA ligation activity.

#### 3. Bead Purification (6min)

1) Add 30 µL nuclease-free $H_2O$ to the 10 µL sample (final volume: 40 µL).
2) Add 72 µL (1.8X) RNA CleanXP beads (Beckman Coulter) and mix thoroughly by pipetting.
3) Incubate at room temperature for 3 min.
4) Place the tube on a magnetic stand until the solution clears.
5) Discard the supernatant. Wash the beads 3 times with 200 µL freshly prepared 70% ethanol.
6) Air-dry the beads, then elute RNA in 10 µL nuclease-free $H_2O$.
7) Incubate at room temperature for 3 min, place on magnet, and transfer the supernatant to a new tube.

**Note:** RNA CleanXP beads can be substituted with other SPRI-based magnetic beads exhibiting comparable RNA-binding properties, including homemade preparations. Philippe Jolivet, Joseph W. Foley 2020. SPRI bead mix. **protocols.io** https://dx.doi.org/10.17504/protocols.io.bnz4mf8w

##### 4. Reverse Transcription (1hour 20min)

1) Add the following to the purified RNA, incubate at **65 °C for 5 min**, then place immediately on ice.

| Reagent | Volume(uL) |
|---|---|
| Random hexamer (20μM) | 1 |
| dNTPs (10mM) | 1 |
| Sample from Step 3 | 10 |
| total | 12 |

2) Add the following reagents, then run this program: 25 °C, 10 min; 42 °C, 50 min; 70 °C, 15 min.

| Reagent | Volume(uL) |
|---|---|
| 5X First-strand buffer | 4 |
| DTT (0.1M) | 2 |
| Nuclease-free H2O | 1 |
| SuperScript II (200U/μL) **** | 1 |

****SuperScript II (200U/μL) can be substituted with other reverse transcriptase with equivalent activity such as RT-MashUP (Alekseenko et al. Sci Rep. 2021doi: 10.1038/s41598-020-80352-8)

##### 5. RNA Depletion and Beads Purification (30 min)

1) Add 8 μL 100 mM NaOH to each sample and incubate at 65 °C for 20 min to hydrolyze RNA.
2) Neutralize with 8 μL 100 mM Tris-HCl.
3) Add 65 μL (1.8X) AMPure XP beads (Beckman Coulter), mix well, and incubate at room temperature for 3 min.
4) Place the tube on a magnetic stand, wait until clear, discard supernatant.
5) Wash beads 3 times with 200 μL freshly prepared 70% ethanol.
6) Air-dry beads and elute DNA in 10 μL nuclease-free $H_2O$.
7) Incubate at room temperature for 3 min, place on magnet, and transfer the supernatant to a new tube.

**Note:** AMPure XP beads can be substituted with other SPRI-based magnetic beads with comparable DNA-binding properties, including homemade alternatives.

##### 6. PCR Amplification (30 min)

Add the following reagents and run the PCR program below.

| Reagent | Volume(uL) |
|---|---|
| 2X Phusion HSII High-Fidelity master mix***** | 9 |
| PE1 (NEBi5, 10μM) | 0.5 |
| PE2 (PE2_MPX, 10μM) | 0.5 |
| Sample from previous step | 10 |
| total | 20 |

*****2X Phusion HSII High-Fidelity master mix can be substituted with other high-fidelity DNA polymerases with dNTP-containing buffer systems and comparable performance.

PCR Program:

| Temperature | Time | Cycles |
|---|---|---|
| 98ºC | 30 sec | 1 |
| 98ºC | 20 sec | |
| 65ºC | 30 sec | 15X |
| 72ºC | 30 sec | |
| 72 ºC | 7 min | 1 |
| 4 ºC | Hold | ∞ |

##### 7. Library Purification and Size Selection (10 min)

1) Add 80 µL nuclease-free $H_2O$ to bring total volume to 100 µL.
2) Add 60 µL (0.6X) AMPure XP beads, mix thoroughly, and incubate at room temperature for 3 min.
3) Place on magnet, transfer 160 µL supernatant to a new tube containing 20 µL (0.2X) AMPure XP beads, mix, and incubate 3 min.
4) Place on magnet, discard supernatant, and wash beads 3 times with 200 µL freshly prepared 70% ethanol.
5) Air-dry beads and elute DNA in 10 µL nuclease-free $H_2O$.
6) Incubate at room temperature for 3 min, place on magnet, and transfer 10 µL supernatant to a clean tube.
7) Quantify libraries using Qubit and assess size distribution with Bioanalyzer.

##### 8. Sequencing Compatibility

- Illumina: Libraries are directly compatible with Illumina sequencing platforms.
- Nanopore: For Oxford Nanopore sequencing, libraries can be used as template for the the Direct Ligation Kit v14 (SQK-LSK114) following the manufacturer's protocol (~1 hour).

**Table: Time and Estimated Per-Sample Cost of s5PSeq Library Preparation**

| Step | Time (hour) | Cost (USD) |
|---|---|---|
| 1. rRNA Blocking Oligo Hybridization | 0.5 | 0.1 |
| 2. ssRNA Ligation | 1.0 | 0.7 |
| 3. Bead Purification | 0.1 | 1.5 |
| 4. Reverse Transcription | 1.3 | 4.0 |
| 5. RNA Depletion and Beads Purification | 0.4 | 1.3 |
| 6. PCR Amplification | 0.5 | 0.9 |
| 7. Library Purification and Size Selection | 0.2 | 1.5 |
| **Total** | ~4 | ~10 |

**Note:** Total costs are estimated based on commercial reagent pricing at small-scale use (per sample basis). Significant reductions are possible using homemade SPRI bead protocols and open-source enzyme preparations (~2 USD per sample), with comparable performance (data not shown).
