## Supplementary material for "A streamlined nanopore-compatible 5PSeq protocol for rapid phenotypic antimicrobial sensitivity testing": Interactive s5PSeq: bsub.html

bam

fivepseqfivepseq

### bam

### Input and run configurations

##### Main input:

|  |  |
| --- | --- |
| Bam directory: | /cfs/klemming/scratch/h/hinliu2/simplified\_fivepseq/bsub/prefivepseq/bam/\*.bam |
| Bam files: | bsub\_ht5pseq bsub\_s5pseq1 bsub\_s5pseq2 |
| Genome file: | /cfs/klemming/projects/supr/sllstore2017018/nobackup/projects/Hong/genome/Bsub/fa/GCF\_000009045.1\_ASM904v1\_genomic.fna |
| Annotation file: | /cfs/klemming/projects/supr/sllstore2017018/nobackup/projects/Hong/genome/Bsub/gff/GCF\_000009045.1\_ASM904v1\_genomic.gff |

##### Auxiliary files

|  |  |
| --- | --- |
| Gene set file: | None |
| Gene filter file: | None |

##### Other options

|  |  |
| --- | --- |
| Output directory: | /cfs/klemming/scratch/h/hinliu2/simplified\_fivepseq/bsub/fivepseq2/fivepseq\_plots |
| Conflict mode: | add |
| Span size: | 100 |
| Transcript type: | protein\_coding |
| Outlier detection p value [--op]: | 0 |
| Down-sampling constant [--ds] | None |
| Masking transcript boundaries for codon counts [--no-mask --mask\_codon\_size]: | by 20 positions |
||  |
| --- | --- |
| Dipeptide relative counts sorted at A-site relative position [--dipeptide\_pos]: | -14 |
||  |
| --- | --- |
| Tripeptide relative counts sorted at A-site relative position [--tripeptide\_pos]: | -11 |

  |  |

### Summary statistics

| Sample | Library size (M) |
| --- | --- |
| bsub\_ht5pseq | 0.09110 |
| bsub\_s5pseq1 | 0.09991 |
| bsub\_s5pseq2 | 0.06358 |

### Report files

##### Main

|  |  |
| --- | --- |
| Main report | main/bam\_main.html |
| Combined samples | main/bam\_combined.html |

##### Supplement

|  |  |
| --- | --- |
| Amino acid line-charts | supplement/bam\_amino\_acid\_linecharts.html |
| Codon line-charts: | supplement/bam\_codon\_linecharts.html |
| Codon heatmaps: | supplement/bam\_codon\_heatmaps.html |
| Dipeptide linecharts: | supplement/bam\_dipeptide\_linecharts.html |
| Tripeptide linecharts: | supplement/bam\_tripeptide\_linecharts.html |
| Dicodon linecharts: | supplement/bam\_dicodon\_linecharts.html |
| Tricodon linecharts: | supplement/bam\_tricodon\_linecharts.html |

##### Comparisons

|  |  |
| --- | --- |
| Differential heatmaps: | comparison/bam\_differential\_heatmaps.html |

##### Gene sets

|  |  |
| --- | --- |
| Compare samples in each gene-set | None |
| Compare gene-sets in each sample | None |

---

These plots are generated with fivepseq version 1.3.5.
For more information on usage and citation, please visit 
our homepage .
